# CircRNA_33702 promotes renal fibrosis by targeting the miR-29b-3p/WISP1 pathway

**DOI:** 10.1101/2022.02.28.482440

**Authors:** Kai Ai, Lei Yi, Yinhuai Wang

## Abstract

Growing evidence suggest that circular RNAs (circRNAs) are critical mediators in renal diseases. However, there have been very few reports about the role of circRNAs in renal fibrosis. In this study, circRNA_33702 was found to be upregulated, both in UUO mice and in TGF-β 1-treated BUMPT cells. Whilst knockdown of circRNA_33702 was shown to relieve the TGF-β 1-induced expression of collagen I, collagen III and fibronectin; overexpression of circRNA_33702 was found to exert an inhibitory effect on the expression of the same genes. Mechanistically, circRNA_33702 was demonstrated to bind directly with miR-29b-3p and inhibit its expression. Moreover, WISP1 was identified as a target of miR-29b-3p and the expression of WISP1 was observed to be repressed by miR-29b-3p. Notably, knockdown of circRNA_33702 was found to attenuate the expression of collagen I, collagen III and fibronectin by inhibiting the expression of WISP1, and the observed inhibitory effect can be reversed by miR-29b-3p inhibitor. Finally, inhibition of circRNA_33702 was shown to attenuate interstitial fibrosis in UUO mice via the miR-29b-3p/WISP1 axis. In general, our data show that circRNA_33702 may promote renal fibrosis via the miR-29b-3p/WISP1 axis, which may potentially be developed as a new therapeutic target.

## Introduction

As the final pathway for all chronic kidney diseases, renal tubulointerstitial fibrosis (TIF) can lead to renal failure resulting from the deposition of extracellular matrix (ECM) and from tubular cell loss.(1–3) Renal fibrogenesis involves most kidney cell types, with tubular epithelial cell being the key target in a variety of kidney injuries.(3, 4) The pathogenesis of TIF involves multiple molecules, such as TGF-β1 that has been widely accepted to play a key role in TIF, and WNT1-inducible signaling pathway protein-1 (WISP-1) that has been suggested as a mediator of TGF-β1-induced renal fibrosis in tubular epithelial cells.(5–8) However, the pathogenetic mechanism of TIF induced by the TGF-β1/WISP-1 pathway remains unclear.

Circular RNAs (circRNAs), a novel class of non-coding RNA is stable against exoribonuclease due to its closed continuous loop that is covalently linked.(9) CircRNAs play important roles in transcription and protein translation. The main function of circRNAs is to cushion microRNAs (miRNAs), which means that circRNAs can interfere miRNAs and their target genes that participate in the progression of various diseases.(10) There have been a number of studies that demonstrate circRNAs as important regulators in multiple diseases, especially in cancers.(11–13) Many recent studies have focused on the link between circRNAs and several renal diseases like acute renal injury, chronic nephritis and diabetic nephropathy.(14, 15)

However, the function of circRNAs on renal fibrosis remains largely unknown. In this study, we found that circRNA_33702 was upregulated in both UUO mice and in TGF-β1-treated BUMPT cells. We then explored the effect of circRNA_33702 on renal fibrosis in BUMPT cells. Moreover, the downstream target of circRNA_33702 and the underlying mechanism of circRNA_33702 in renal fibrosis were further investigated. Our results may facilitate the development of a novel therapeutic target for renal fibrosis.

## Materials and Methods

### Antibodies and reagents

Anti-collagen I, anti-collagen III, anti-fibronectin and anti-WISP1 antibodies were obtained from Abcam (Cambridge Science Park, Cambridge, UK), whereas anti-GAPDH antibody was purchased from Proteintech North America (Rosemont, IL, USA). Luciferase Assay kit was obtained from BioVision (Milpitas, CA, USA).

### Cell culture and treatments

BUMPT cells were cultured in high glucose DMEM (Thermo Fisher Scientific) supplemented with 10% fetal bovine serum (FBS) and penicillin-streptomycin (100 U/ml and 100 μg/ml, respectively) at 37°C with CO_2_. The cells were transfected with miR-29b-3p inhibitor (100 nM), miR-29b-3p mimic (100 nM), circRNA_33702 siRNA (100 nM), circRNA_33702 plasmid, WISP1 siRNA (100 nM), or negative control (Ruibo, Guangzhou, China) using Lipofectamine 2000 (Life Technologies, Carlsbad, CA, USA). Twenty-four hours after transfection, the cells were treated with serum-free medium overnight, followed by treatment with saline or TGF-β1 (5 ng/ml) for 24 hours.

### Luciferase reporter assays

Luciferase vector constructs containing full length circRNA_33702 (WT-Luc-circRNA_33702), WISP1 3’ UTR (WT-Luc-WISP1), circRNA_33702 fragment that interacts with miR-29b-3p, WISP1 fragment that interacts with miR-29b-3p, mutated full length circRNA_33702 lacking the miR-29b-3p-interacting fragment (MUT-Luc-circ _33702) and mutated WISP1 3’ UTR lacking the miR-29b-3p-interacting fragment (MUT-Luc-WISP1), respectively were used in this study. Specific target activity was calculated based on the relative luciferase activity ratio of firefly to renilla. All plasmids were obtained from Vigene Biosciences (Jinan, shangdong, China).

BUMPT cells were co-transfected with WT-Luc-circRNA_33702 or MUT-Luc-circRNA_33702, with or without miR-29b-3p mimics. Luciferase reporter assay was performed after incubation for 48h as previously described.(16–18) Finally, firefly and renilla luciferase activities were measured using Dual-Glo Luciferase Assay System (Promega, Madison, WI, USA) with SpectraMaxM5 (Molecular Devices, Sunnyvale, CA, USA).

### Animal models

UUO model was established in C57BL/6 mice (male, aged 10–12 weeks) as previously described by ligating the left ureter with injection of sodium pentobarbital, using sham-operated mice as controls. The mice were injected twice a week with circ_33702 siRNA(15 mg/kg) or saline via tail vein. The mice were housed on a 12-h light/12-h dark cycle with free access to food and water. All animal studies were performed according to the institutional guidelines of Committee for the Care and Use of Laboratory Animals of Second Xiangya Hospital, People’s Republic of China.

### Histology and immunohistochemistry (IHC)

The histology of harvested kidney tissues was analyzed using hematoxylin-eosin (H&E) staining. Fibrotic area was assessed by masson’s trichrome staining. IHC analysis was performed using anti-collagen I (1:100 dilution), anti-collagen III (1:100 dilution) and anti-FN (1:100 dilution) antibodies as previously described.(19) IHC images were analyzed using an Olympus microscope equipped with UV epi-illumination.

### Real-time qPCR

Total RNA was isolated from BUMPT cells and mouse kidney tissues using Trizol reagent (Invitrogen, Carlsbad, CA, USA) according to the manufacturer’s protocol., Approximately 40 ng of total RNA was used to synthesize cDNAs using M-MLV Reverse Transcriptase (Invitrogen). To quantify the expression levels of circRNA, mRNA and miRNA, real-time qPCR was performed using Bio-Rad (Hercules, CA, USA) IQ SYBR Green Supermix with Opticon (MJ Research, Waltham, MA, USA) according to the manufacturer’s instructions. The nucleotide sequence of miR-29b-3p was retrieved from GenBank database (gen ID:387223). The nucleotide sequence of circRNA_33702 was listed in supplementary data. The sequences of primers used in this study are as follows: 5’-CCAGCTCTTCTGAAGGAAGCACAG-3’ (circRNA_33702-forward), 5’-TGGCTTAAGGTCCTCCTCAGGTTC-3’ (circRNA_33702-reverse), 5’-CGCGTAGCACCATTTGAAATC-3’ (miR-29b-3p-forward), 5’-AGTGCAGGGTCCGAGGTAT-3’ (miR-29b-3p-reverse), 5’-GGTCTCCTCTGACTTCACA-3’ (GAPDH-forward) and 5’-GTGAGGGTCTCTCTCTTCCT-3’ (GAPDH-reverse). The sequences for U6 primers were used as described previously.(20) Relative expression of target genes normalized against GAPDH or U6 was calculated using the 2^−ΔΔCt^ method.

### Western blot analysis

Total protein from BUMPT cells and mouse kidney tissues was extracted by cell lysis, followed by centrifugation at 12,000r at 4°C for 15 min. The resulting supernatant fraction of protein lysates was mixed with SDS-PAGE sample buffer containing β-ME before being boiled at 100°C for 5 min. An equal amount of protein lysates was separated by SDS-PAGE before being transferred onto PVDF membrane (Amersham, Buckinghamshire, UK) for 120 minutes at 290mA.(19, 21, 22) Transferred membranes were subsequently incubated respectively with primary antibodies against collagen I, collagen III, fibronectin, WISP1 and GAPDH overnight at 4°C, followed by incubation with secondary antibody at room temperature for 60 minutes. ECL kit (Millipore) was used to display target protein bands.

### Fluorescence in site hybridization (FISH)

The nucleotide sequence of circRNA_33702 and miR-29b-3p was hybridized using fluorescent probes (Ruibo, Guangzhou, China) as previously described.(23) Briefly, BUMPT cells and mouse kidney tissues were incubated with fluorescent probes in hybridization buffer overnight at 4°C. After a washing step with SSC buffer, the cell nuclei were stained with DAPI while 18S rRNA was being used as a cytoplasmic control. Cell images were visualized using a laser scanning confocal microscope.

### Statistical analyses

Statistical analyses were performed using Graphpad Prism 7. Difference between two groups were compared using two-tailed Student’s t tests. Difference between multiple groups were compared using one-way ANOVA. Quantitative data were presented as means ± standard deviation (SD). P< 0.05 was considered statistically significant for all statistical comparisons.

## Results

### The expression of circRNA_33702 was up-regulated in the progression of renal fibrosis

Our previous study has shown significantly upregulated circRNAs in the UUO-induced renal fibrosis model mice kidney tissues at days 3 and 7 after surgery compared to those from sham controls on days 3 and 7 by circRNA microarray analysis. CircRNA_33702 was one of the most highly expressed circRNA in the kidney tissues and showed a 8.92- and 15.55-fold higher expression in the UUO group on days 3 and 7, respectively, compared to the sham group.(24) To investigate the expression site of circRNA_33702, BUMPT cells were treated with TGF-β1 before being subjected to FISH analysis. The cell nuclei were stained by DAPI, while 18S rRNA (cytoplasmic positive) and circRNA_33702 were labeled with CY3. As shown in Figure 1A, circRNA_33702 was found to be localized in the cytoplasm of BUMPT cells. To evaluate the expression profile of circRNA_33702, real-time qPCR results showed that circRNA_33702 was induced both in TGF-β1-treated BUMPT cells and in UUO mice kidneys (Figure 1B&1C), indicating that circRNA_33702 may be involved in renal fibrosis.

**Figure 1.**
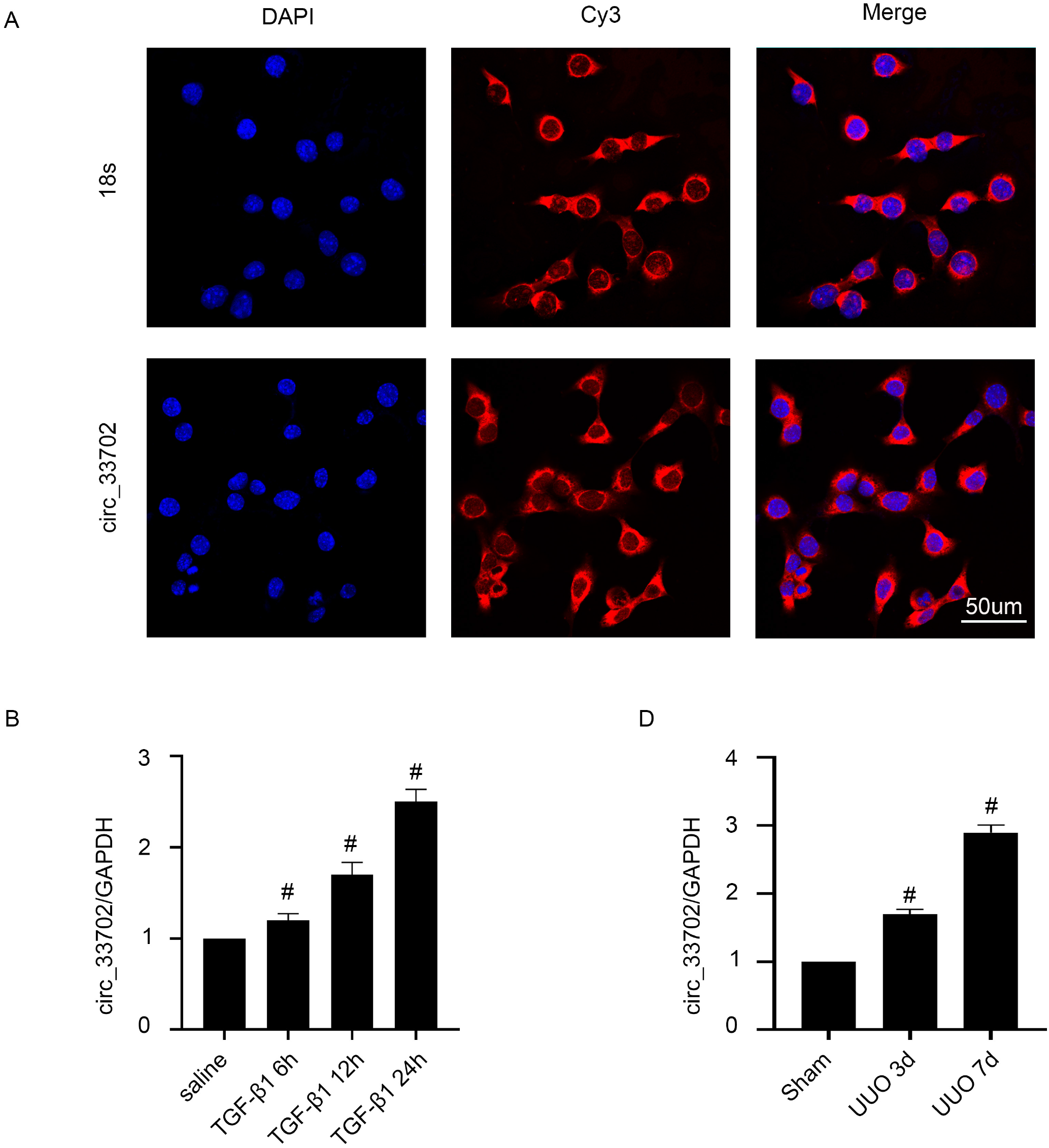
CircRNA_33702 was induced both in TGF-β1-treated BUMPT cells and in UUO mouse kidneys. Male C57BL/6 mice were treated with UUO for 0-7 days, while BUMPT cells were treated with 5 ng/ml TGF-β1 for 0-24h. (A) Intracellular localization of circRNA_33702 in BUMPT cells. (B&C) Real-time qPCR analyses of circRNA_33702 expression in TGF-β1-treated BUMPT cells and in UUO mouse kidneys. Data are expressed as mean ± SD (n = 6). #p < 0.05, versus saline or sham group.

### Inhibition of circRNA_33702 expression attenuated the TGF-β1-induced expression of collagen I, collagen III and fibronectin

Next, we examined the role of circRNA_33702 in TGF-β1-treated BUMPT cells. As shown in Figure 2A, transfection of BUMPT cells with circRNA_33702 siRNA inhibited the TGF-β1-induced expression of circRNA_33702. Furthermore, inhibition of circRNA_33702 expression attenuated the expression of collagen I, collagen III and fibronectin, both at basal levels and TGF-β 1-induced levels (Figure 2B-2E). These results indicated that inhibition of circRNA_33702 is capable of relieving TGF-β1-induced renal fibrosis in BUMPT cells, suggesting that circRNA_33702 may play a role as a pro-fibrosis mediator in BUMPT cells.

**Figure 2.**
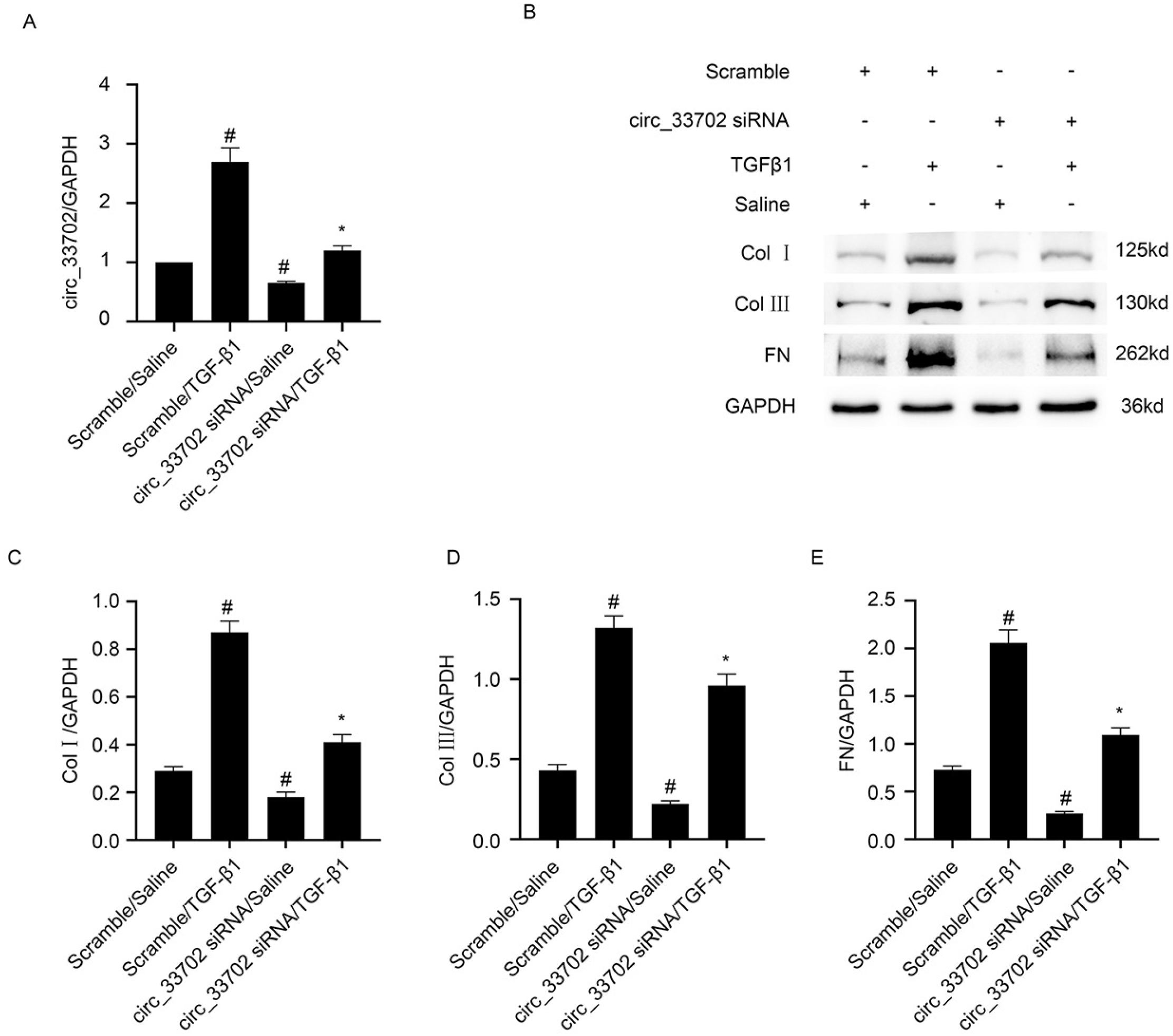
Inhibition of circRNA_33702 relieved the TGF-β1-induced expression of collagen I, collagen III, and fibronectin. BUMPT cells were transfected with 50 nM circRNA_33702 or scrambled siRNA before being treated with TGF-β1 or saline for 24 h. (A) Expression of circRNA_33702 detected by real-time qPCR. (B) Protein expression of collagen I, collagen III and fibronectin detected by western blotting. (C-E) Densitometric measurement of western blot bands for collagen I (C), collagen III (D) and fibronectin (E). Data are expressed as mean ± SD (n = 6). #p < 0.05, versus control group. *p < 0.05, versus TGF-β1 group.

### Overexpression of circRNA_33702 aggravated the TGF-β1-induced expression of collagen I, collagen III, and fibronectin

To further explore the effect of circRNA_33702 on TGF-β 1-induced renal fibrosis, BUMPT cells were transfected with circRNA_33702 plasmid constructs. As shown in Figure 3A, the expression of circRNA_33702 in BUMPT cells was increased after transfection of circRNA_33702. The overexpression of circRNA_33702 enhanced the expression of collagen I, collagen III and fibronectin, not only at basal levels, but also at TGF-β 1-induced levels (Figure 3B-3E). These results further demonstrate the pro-fibrosis function of circRNA_33702 in TGF-β1-treated BUMPT cells.

**Figure 3.**
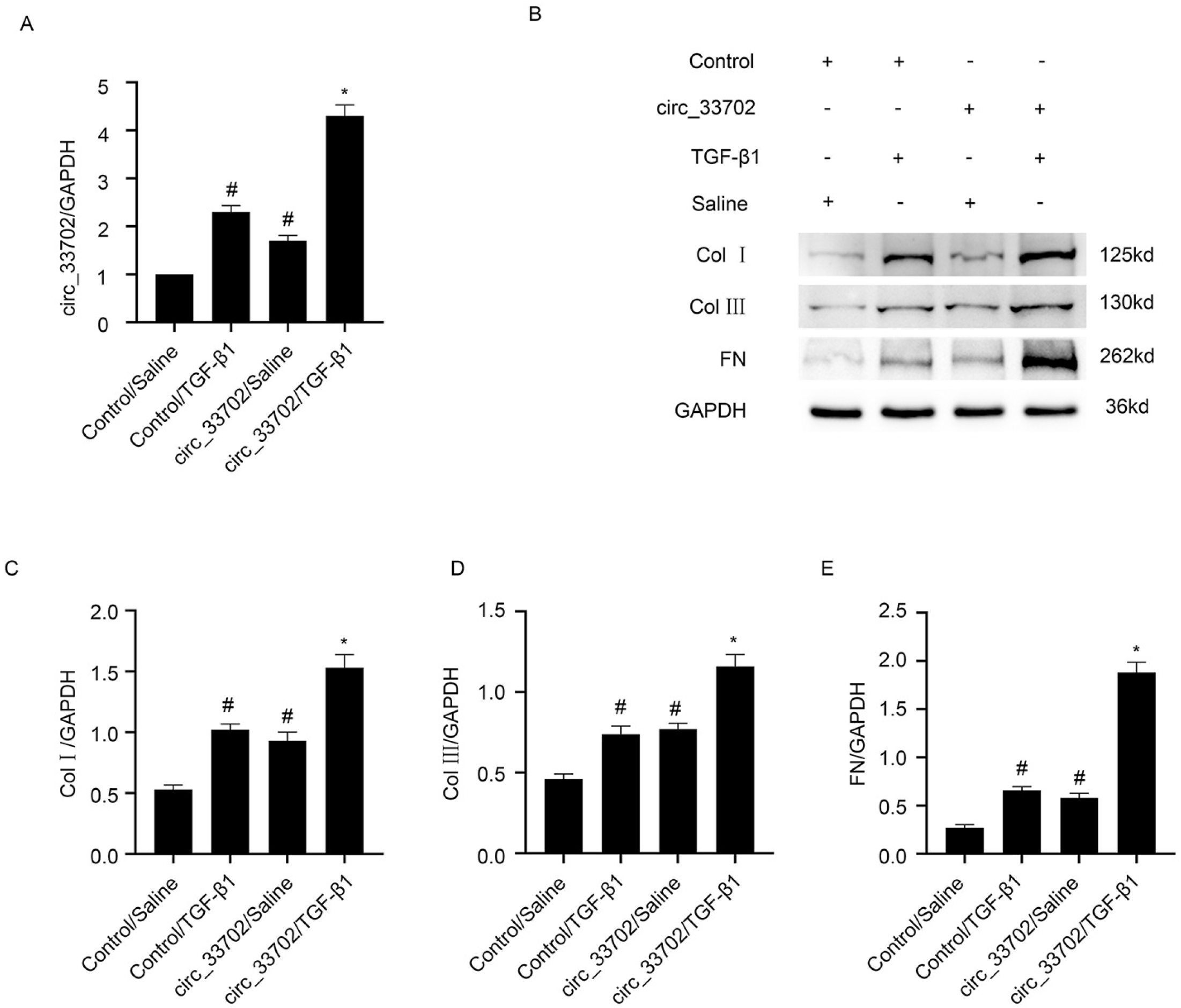
Overexpression of circRNA_33702 aggravated TGF-β 1-induced expression of collagen I, collagen III, and fibronectin. BUMPT cells were transfected with circRNA_33702 plasmid construct or control plasmid before being treated with TGF-β1 or saline for 24 h. (A) Expression of circRNA_33702 detected by real-time qPCR. (B) Protein expression of collagen I, collagen III and fibronectin detected by western blotting. (C-E) Densitometric measurement of western blot bands for collagen I (C), collagen III (D) and fibronectin (E). Data are expressed as mean ± SD (n = 6). #p < 0.05, versus control group. *p < 0.05, versus TGF-β1 group.

### CircRNA_33702 binds directly to miR-29b-3p

Prediction using RegRNA 2.0 software showed that circRNA_33702 exhibits two potential binding sites of miR-29b-3p, indicating that miR-29b-3p may be a potential target of circRNA_33702 (Figure 4A). Luciferase reporter assays demonstrated that miR-29b-3p mimic could inhibit the luciferase activity of circRNA_33702 WT and circRNA_33702-MUT1, but not circRNA_33702-MUT2, implying that the fragment at base position 741-747 of circRNA_33702 is the real potential binding site of miR-29b-3p (Figure 4B). Intracellular co-localization analysis showed overlapping signals of circRNA_33702 and miR-29b-3p in the cytoplasm of BUMPT cells and renal tubular cells of C57BL/6 mouse kidneys, indicating that circRNA_33702 and miR-29b-3p may have physical interaction (Figure 4C). The TGF-β1-supressed expression of miR-29b-3p was enhanced through knockdown of circRNA_33702, and this effect was reversed through overexpression of circRNA_33702 (Figure 4D & 4E). These results demonstrated that circRNA_33702 can directly interact with miR-29b-3p.

**Figure 4.**
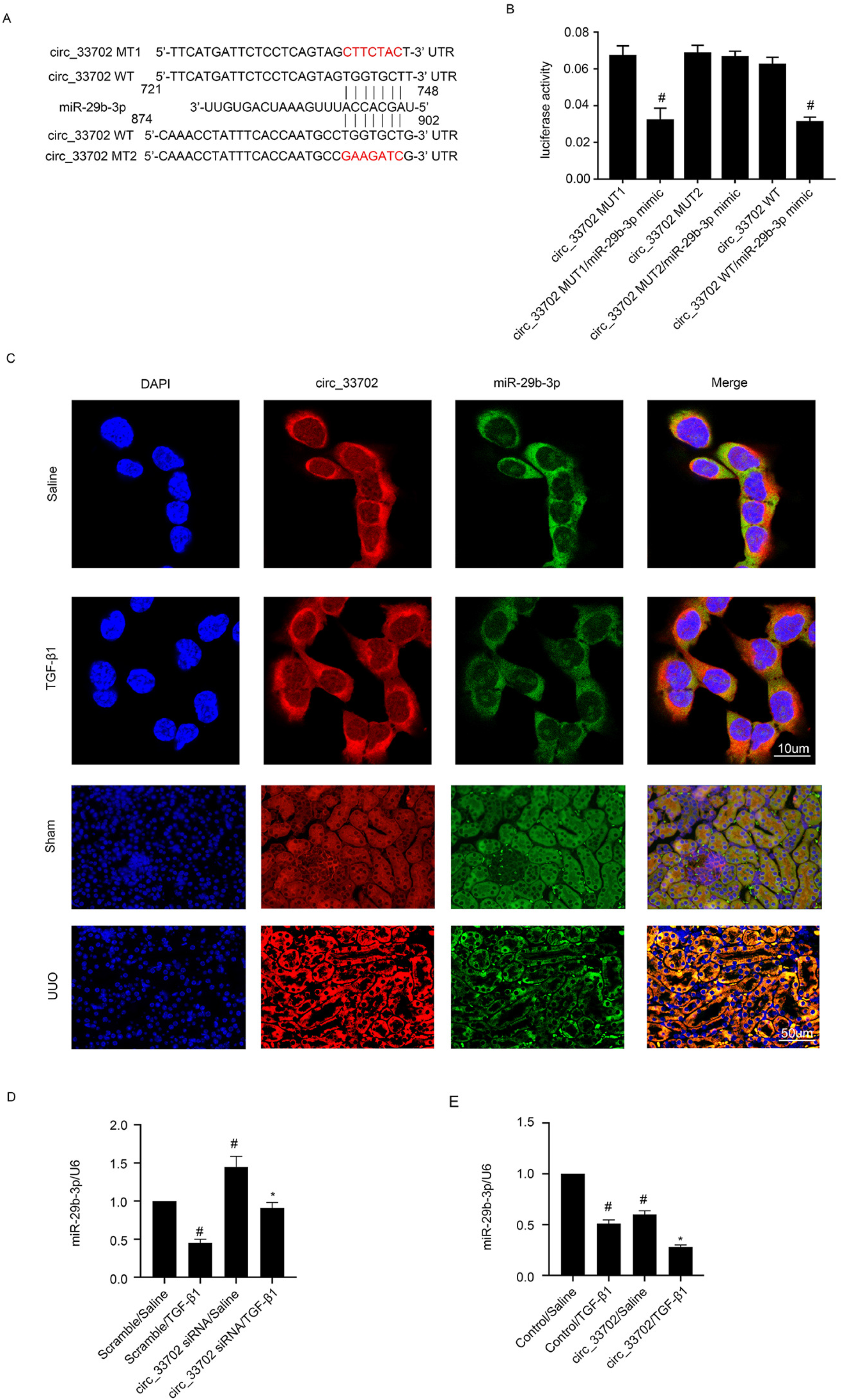
Negative correlation between circRNA_33702 and miR-29b-3p. (A) Complementary strand of circRNA_33702 to miR-29b-3p predicted by RegRNA 2.0 software. (B) Luciferase activity measurement of cells after co-transfection with circRNA_33702-WT or circRNA_33702-MUTs and miR-29b-3p or scrambled siRNA. (C) Intracellular co-localization of circRNA_33702 and miR-29b-3p in BUMPT cells and in the kidneys of C57BL/6 mice. (D and E) Expression of miR-29b-3p detected by real-time qPCR. Data are expressed as mean ± SD (n = 6). #p< 0.05, versus scrambled siRNA or control with saline group; *p< 0.05, siRNA circRNA_33702 or circRNA_33702 with TGF-β1 group versus scrambled siRNA or control with TGF-β1 group or circRNA_33702 MUT1/miR-29b-3p mimic or circRNA_33702 WT/miR-29b-3p mimic versus other groups.

### MiR-29b-3p inhibited TGF-β1-induced ECM accumulation in BUMPT cells

Previous studies have suggested that miR-29b-3p has an anti-fibrosis role in liver and cardiac fibrosis(25–27), but the role of miR-29b-3p in renal fibrosis remains unclear. Real-time qPCR confirmed that miR-29b-3p mimic increased the expression of miR-29b-3p in BUMPT cells (Figure 5A). Following transfection of BUMPT cells with miR-29b-3p mimic, the TGF-β1-induced expression of collagen I, collagen III and fibronectin was decreased (Figure 5B-5E). These results showed that miR-29b-3p inhibits TGF-β1-induced ECM accumulation in BUMPT cells.

**Figure 5.**
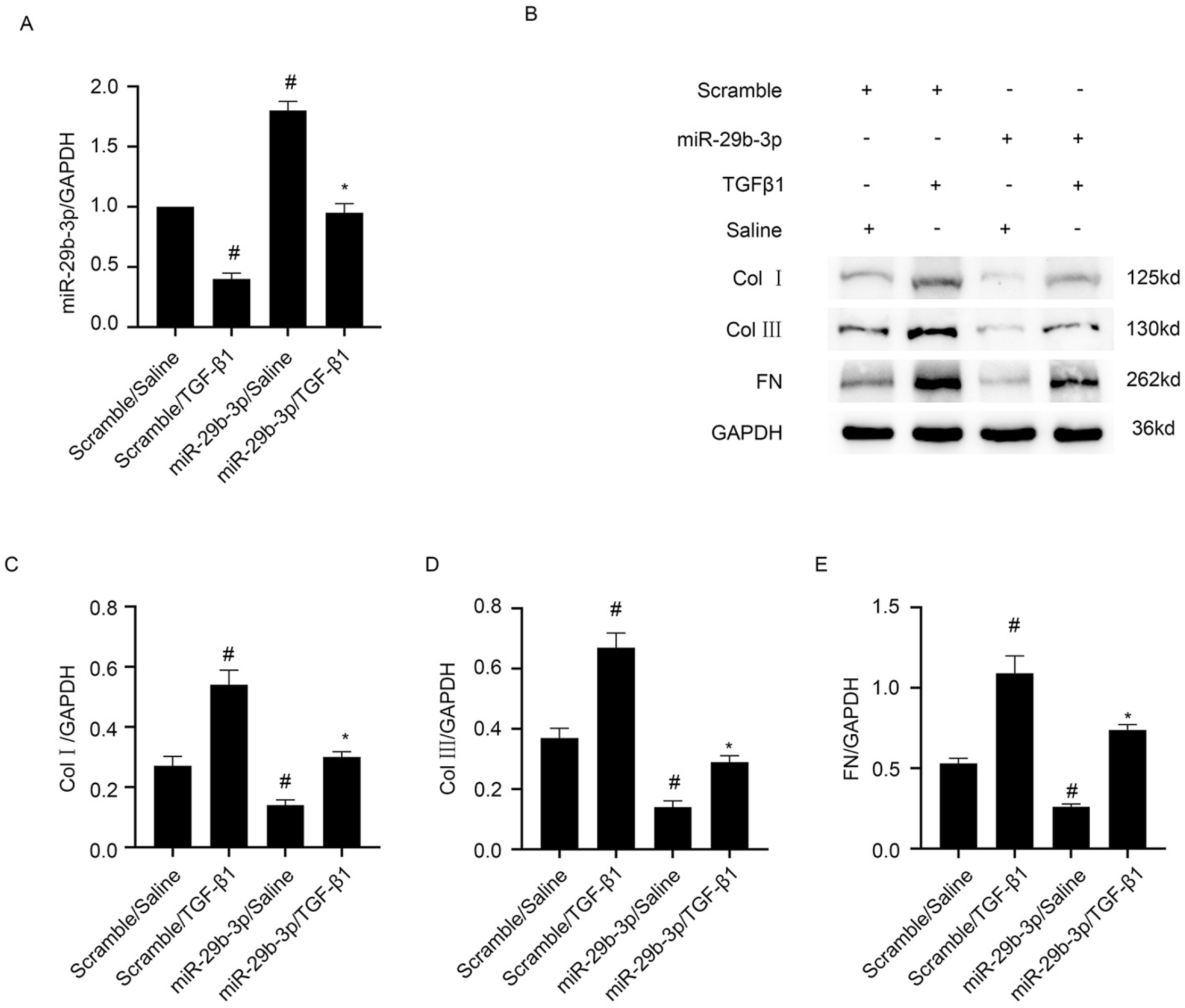
Overexpression of miR-29b-3p relieved the TGF-β1-induced expression of collagen I, collagen III, and fibronectin. BUMPT cells were transfected with 100 nM miR-29b-3p or scrambled siRNA and before being treated with TGF-β1 or saline for 24 h. (A) Real-time qPCR analysis of miR-29b-3p expression. (B) Western blot analysis of collagen I, collagen III and fibronectin expression. (C-E) Densitometric measurement of western blot bands for collagen I (C), collagen III (D) and fibronectin (E). Data are expressed as mean ± SD (n = 6). #p < 0.05, versus scrambled siRNA group. *p < 0.05, versus TGF-β1 group.

### WISP1 is a target gene of miR-29b-3p

WISP1 has been suggested to be a mediator of TGF-β1-induced renal fibrosis. As shown in Figure 6A, WISP1 was predicted by miRbase software as a potential target of miR-29b-3p. Luciferase assay demonstrated that, whilst miR-29b-3p mimic inhibited the luciferase activity of WISP1-WT, it had no effect on the luciferase activity of WISP1-MUT (Figure 6B). Furthermore, miR-29b-3p mimic was observed to downregulate the TGF-β1-induced expression of WISP1 (Figure 6C-6E). Taken together, WISP1 may be a direct target of miR-29b-3p.

**Figure 6.**
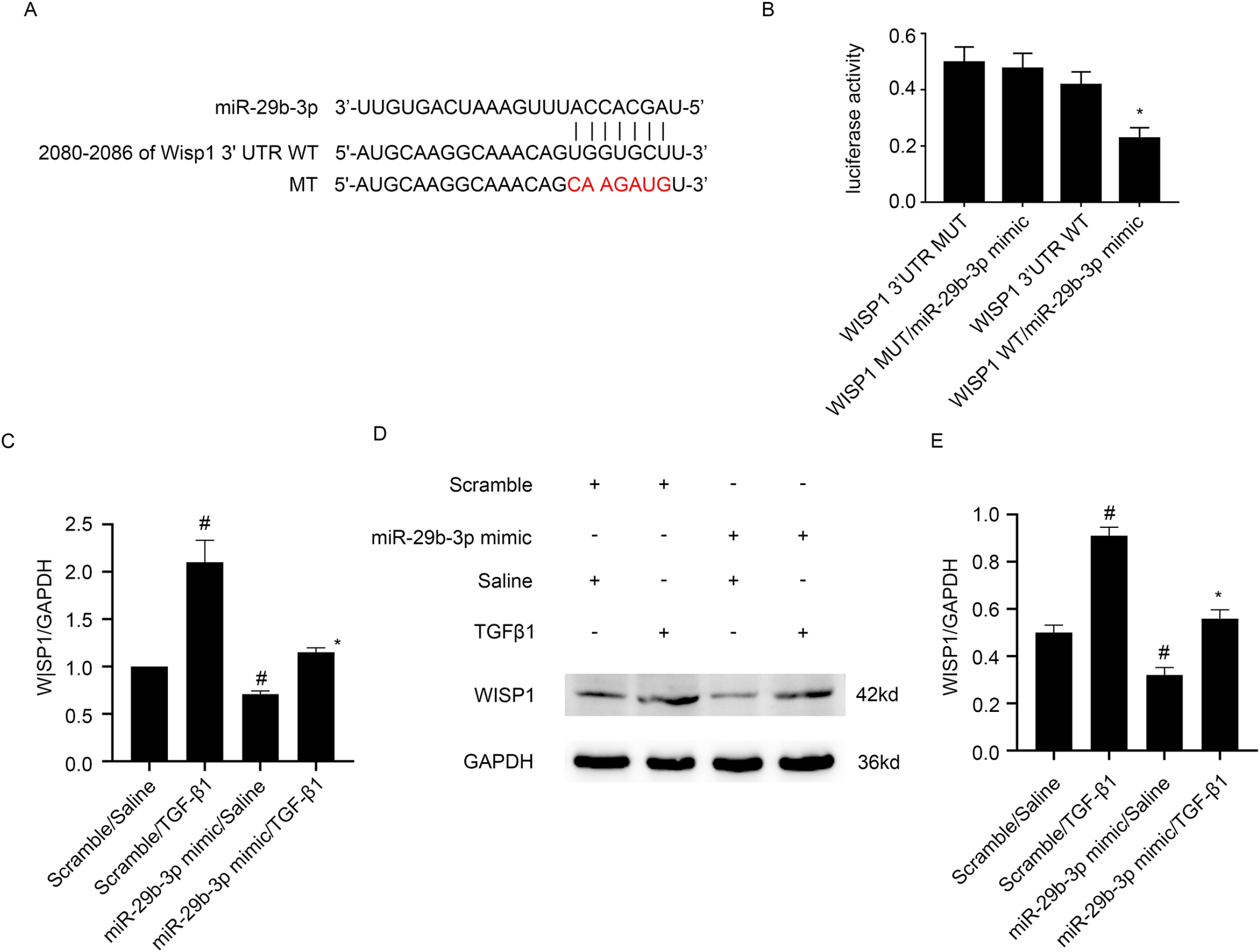
WISP1 was identified as a target gene of miR-29b-3p. BUMPT cells were transfected with miR-29b-3p analog (100 nM) before being treated with TGF-β1 for 24 h. (A) MiR-29b-3p complementary binding sites in the 3’UTR of mice WISP1 mRNA. (B) Analysis of luciferase activities in cells after co-transfection with 3’ UTR luciferase reporter vector of mice WT or MUT-WISP1 and miR-29b-3p or miR-NC. (C) Real-time qPCR analysis of WISP1 mRNA expression. (D) Protein expression of WISP1 detected by western blotting. (E) Densitometric measurement of western blot band for WISP1. #p< 0.05, versus scrambled siRNA group; *p < 0.05, miR-29b-3p analog with TGF-β1 group versus scrambled siRNA with TGF-β1 group or WISP1 WT/miR-29b-3p analog versus other groups.

### MiR-29b-3p reversed the pro-fibrotic effect of circRNA_33702

Next, we explored whether miR-29b-3p is capable of reversing the effect of circRNA_33702 in BUMPT cells. The transfection efficiency of circRNA_33702 siRNA and miR-29b-3p inhibitor were firstly verified by real-time qPCR (Figure 7A&7B). Western blot analysis showed that inhibition of circRNA_33702 expression relieved the TGF-β1-induced expression of collagen I, collagen III, fibronectin and WISP1; while miR-29b-3p inhibitor reversed the effect of circRNA_33702 and increased the expression of collagen I, collagen III, fibronectin and WISP1 (Figure 7C-7G). These observations indicated that circRNA_33702 may promote renal fibrosis by inhibiting the expression of miR-29b-3p.

**Figure 7.**
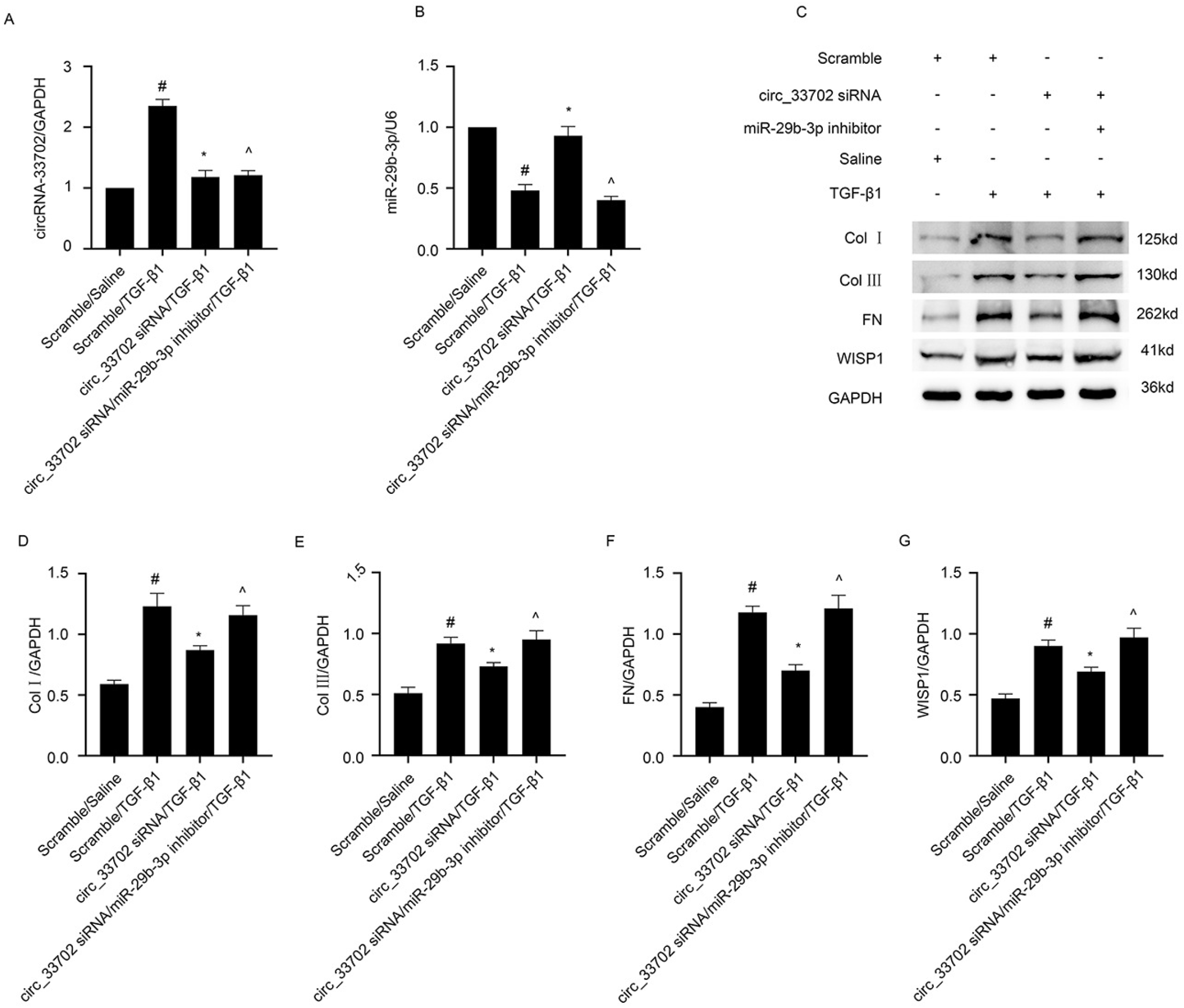
CircRNA_33702 siRNA relieved TGF-β1-induced ECM accumulation, which can be reversed by miR-29b-3p inhibitor. BUMPT cells were co-transfected with circRNA_33702 siRNA and miR-29b-3p inhibitor or scrambled siRNA before being treated with TGF-β1 for 24 h. (A) Expression of circRNA_33702 detected by real-time qPCR. (B) Expression of miR-29b-3p detected by real-time qPCR. Data are expressed as mean ± SD (n = 6). #p< 0.05, scrambled siRNA with TGF-β1 group versus scramble group, *p< 0.05, circRNA_33702 siRNA with TGF-β1 group versus scrambled siRNA with TGF-β1 group; ^p< 0.05, circRNA_33702 siRNA plus miR-29b-3p inhibitor with TGF-β1 group versus circRNA_33702 siRNA with TGF-β1 group.

### Inhibition of circRNA_33702 attenuated UUO-induced renal fibrosis

In order to verify our *in vitro* findings, the expression of circRNA_33702 in male C57BL/6 mice that were on a 7-day UUO treatment was knocked-down by injecting circRNA_33702 siRNA on day 1 before surgery and on day 3 after surgery. As shown in Figure 9A, real-time qPCR indicated that the expression of circRNA_33702 was inhibited, both in UUO mice and in sham control mice, confirming that the knockdown of circRNA_33702 was efficient. The expression of miR-29b-3p that was suppressed in obstructive kidneys was observed to be resumed through the knockdown of circRNA_33702 (Figure 9B). HE staining showed that circRNA_33702 was capable of relieving UUO-induced tubular dilation and atrophy in mouse kidneys; while Masson’s staining demonstrated that circRNA_33702 was effective in attenuating UUO-induced ECM accumulation (Figure 8B). Quantitative analysis of fibrotic areas was performed according to the intensities of HE and Masson’s staining (Figure 8F). Immunohistochemical staining demonstrated that knockdown of circRNA_33702 successfully attenuated the UUO-induced expression of collagen I, collagen III and fibronectin (Figure 8C-8E&8G). Immunoblotting further verified that the UUO-induced expression of collagen I, collagen III, fibronectin and WISP1 were relieved through the knockdown of circRNA_33702 (Figure 9C-9G). Taken together, these data demonstrated that circRNA_33702 may promote renal fibrosis via the miR-29b-3p/WISP1 axis.

**Figure 8.**
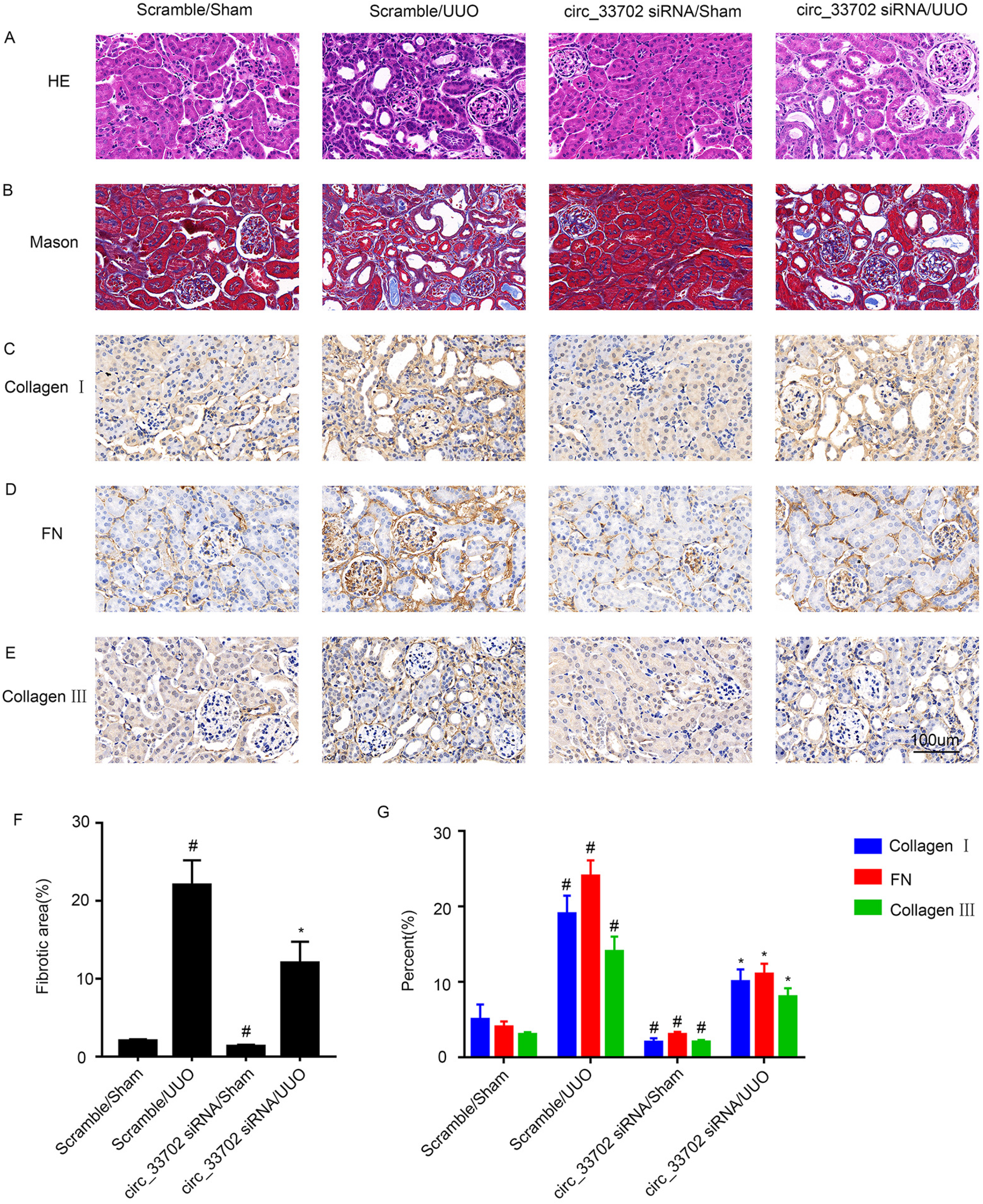
Inhibition of circRNA_33702 attenuated UUO-induced renal fibrosis. The left ureter of male C57BL/6 mice was ligated to establish the UUO model for 7 days. (A) Renal tissues stained with hematoxylin and eosin. (B) Renal tissues stained with Masson’s trichrome. (C-E) Immunohistochemistry of collagen I, collagen III and fibronectin. (F) Quantify tubulointerstitial fibrosis in the kidney cortex. (G) Quantification of immunohistochemistry staining. Data are expressed as mean ± SD (n= 6). #P< 0.05 versus sham group. *P< 0.05 versus scrambled siRNA with UUO group. Original magnification, ×200.

**Figure 9.**
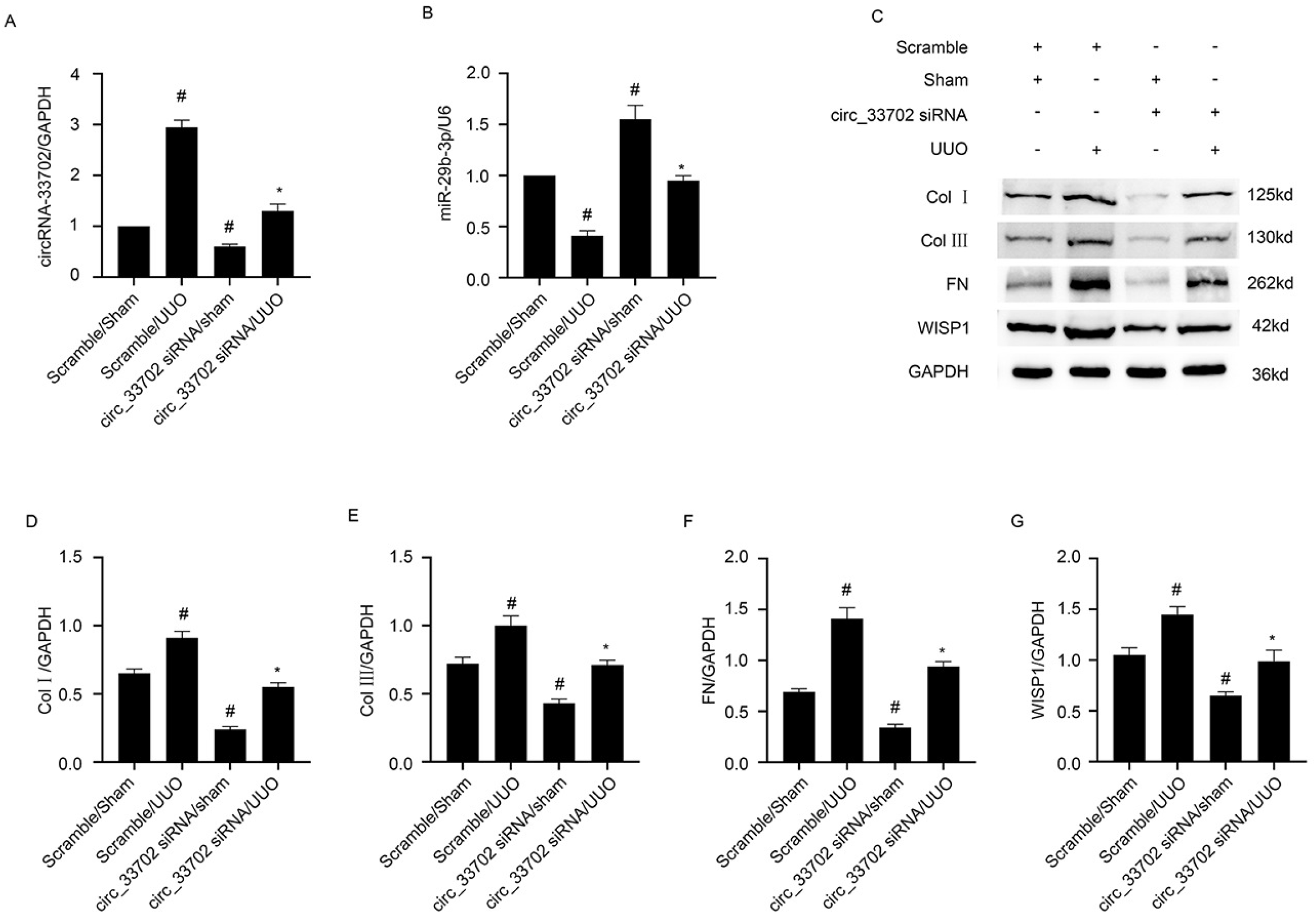
Inhibition of circRNA_33702 attenuated the UUO-induced expression of collagen I, collagen III, fibronectin and WISP1. The left ureter of male C57BL/6 mice was ligated to establish the UUO model for 7 days. (A) Real-time qPCR analysis of circRNA_33702. (B) Real-time qPCR analysis of miR-29b-3p. (C) Protein expression of collagen I, collagen III, fibronectin and WISP1 detected by western blotting. (D-G) Densitometric measurement of western blot bands for collagen I (D), collagen III (E), fibronectin (F) and WISP1(G). #P< 0.05 versus sham group. * P< 0.05 versus scrambled siRNA with UUO group.

## Discussion

With an increasing prevalence globally, renal fibrosis is commonly diagnosed in all chronic kidney diseases. Unfortunately, there are very few available therapeutic options. For the first time, this study identified the role of circRNA_33702 in TGF-β1-treated BUMPT cells and in the obstructive kidneys of mice. Whilst the expression of circRNA_33702 was found to be enhanced in renal fibrosis, overexpression of circRNA_33702 was observed to aggrevate TGF-β1-induced renal fibrosis. Conversely, knockdown of circRNA_33702 was shown to attenuate TGF-β1-induced renal fibrosis. In terms of regulatory mechanism, our results suggest that the expression of miR-29b-3p is negatively regulated by circRNA_33702, and that WISP1 is a target gene of miR-29b-3p. Moreover, renal fibrosis was found to be attenuated through the inhibition of circRNA_33702 in mice UUO model. Our study provides direct evidence that demonstrate the association between the circRNA_33702/ miR-29b-3p/WISP1 axis and renal fibrosis. The findings in this study may facilitate the development of therapeutic interventions for kidney fibrosis (Figure10).

**Figure 10.**
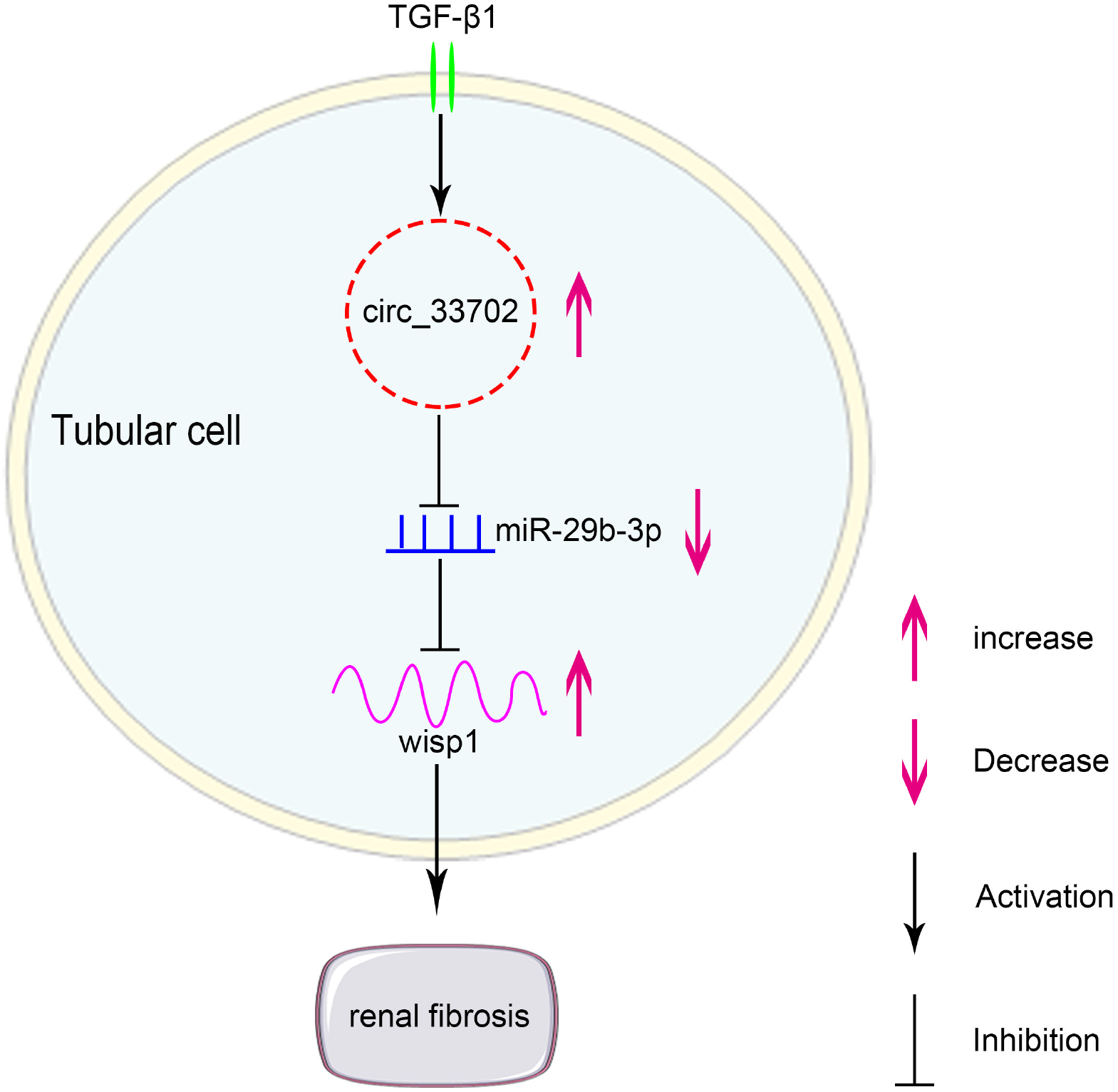
Molecular mechanism of circRNA_33702 in TGF-β1-induced renal fibrosis. The expression of circRNA_33702 was induced in TGF-β1-treated BUMPT cells and in UUO mouse kidneys. CircRNA_33702 sponges miR-29b-3p to upregulate the expression of WISP1, which then promotes renal fibrosis.

An increasing number of studies have demonstrated that circRNAs are involved in renal diseases. Xiong and colleagues have found that the expression of circRNA_ZNF609 is upregulated in renal cell carcinoma, and that the overexpression of circRNA_ZNF609 promotes cell proliferation and invasion via the miR-138-5p/FOXP4 axis.(28) Meanwhile, Chen and colleagues have found that circRNA-LRP6 can upregulate the expression of HMGB1 and activate the TLR4/NF-κB pathway by sponging miR-205, in order to regulate high glucose-induced proliferation, oxidative stress, ECM accumulation and inflammation in mesangial cells.(29) Additionally, Deng and colleagues have demonstrated that circRNA_ANRIL can promote LPS-induced the inflammation and apoptosis of HK2 cells by regulating the miR-9/NF-κB axis.(30) However, there have been very few studies that focus on the effect of circRNA in renal fibrosis. In this study, we found that the expression of circRNA_33702 was up-regulated in TGF-β1-treated BUMPT cells as well as in UUO mice. In addition, whilst circRNA_33702 knockdown was found to aggravate the TGF-β1-induced renal fibrosis; the observed effect was successfully reversed though the overexpression of circRNA_33702. Our data suggest that circRNA_33702 plays a pro-fibrotic role in TGF-β1-treated BUMPT cells.

An increasing number of studies have demonstrated that circRNAs, a new category of endogenous non-coding RNA, can decrease miRNA abundance by sponging miRNA in order to regulate targeted gene expression. In this study, through dual-luciferase reporter assay, circRNA_33702 was demonstrated to be capable of binding miR-29b-3p directly. In addition, through FISH co-localization assay, circRNA_33702 was revealed to interact with miR-29b-3p, both in BUMPT cells and in mouse kidneys. Moreover, miR-29b-3p inhibitor was found to successfully reverse the effect of circRNA_33702 knockdown and subsequently promote ECM accumulation. The observations above demonstrate that miR-29b-3p is a direct target of circRNA_33702.

Many studies have shown that miR-29b-3p is involved in fibrosis. Ni and colleagues have reported that miR-29b-3p, as a direct target of circRNA-HIPK3, can inhibit the proliferation and migration of cardiac fibroblasts, and that the inhibition effect can be altered by targeting α-SMA, COL1A1 and COL3A1.(31) Tao and colleagues have found that miR-29b-3p can prevent granulomatous liver fibrosis through direct inhibition of COL1A1 and COL3A1.(32) Zhang and colleagues have demonstrated that carnosic acid can alleviate bile duct ligation-induced liver fibrosis in rats by modulating the miR-29b-3p/HMGB1/TLR4/ NF-κB signaling pathway.(33) Our study revealed that overexpression of miR-29b-3p can inhibit the TGF-β1-induced ECM accumulation in BUMPT cells, and that WISP1 is a potential target of miR-29b-3p. Furthermore, we demonstrated that miR-29b-3p interacts directly with WISP1, and that miR-29b-3p overexpression can suppress TGF-β1-induced WISP1 expression. Taken together, these results show that WISP1 is a target of miR-29b-3p.

Several studies have shown that WISP1 is involved in the fibrosis of various organs including kidney. Yang and colleagues have explored the functional role and mechanism of WISP1 in renal fibrosis using TGF-β1-treated tubular epithelial cells and UUO mouse models; they have found that WISP1 can upregulate the expression of collagen I, FN and α-SMA, indicating that WISP1 can promote renal fibrosis by inducing autophagy.(7) Chen and colleagues have reported that knockdown of WISP1 can suppress the activation of the wnt/β-actin catenin signaling pathway, which then decreases the EMT of renal TECs in uremic rats.(34) In this study, we demonstrated that knockdown of circRNA_33702 can relieve TGF-β1-induced renal fibrosis through the downregulation of WISP1, and that the observed effect can be reversed by miR-29b-3p inhibitor. Our *in vitro* observations were confirmed using UUO mouse kidney samples, by showing that the inhibition of circRNA_33702 can attenuate renal fibrosis in UUO mice by targeting the miR-29b-3p/WISP1 axis.

In summary, our work demonstrated that the expression of circRNA_33702 is induced in renal fibrosis. In addition, overexpression of circRNA_33702 was found to aggravate TGF-β1-induced renal fibrosis, while knockdown of circRNA_33702 was observed to relieve TGF-β1-induced renal fibrosis. Furthermore, we showed that overexpressed circRNA_33702 directly interacts with miR-29b-3p to upregulate WISP1 expression, which ultimately promotes renal fibrosis. Our findings suggested that circRNA_33702 plays a pro-fibrosis role, and that circRNA_33702 may be a novel therapeutic target of renal fibrosis.

## Acknowledgements

We appreciate all the participants who provide supports for the study.

## Author Contributions

KA and LY collected the data and wrote the manuscript; YW were responsible for the study design.

## Availability of supporting data

All data used during the current study available from the corresponding author on reasonable request.

## Abbreviation

BUMPT: Boston university mouse proximal tubule
UUO: unilateral ureteral obstruction
WISP1: WNT1 inducible signaling pathway protein 1

